# E93 expression and links to the juvenile hormone in hemipteran mealybugs with insights on female neoteny

**DOI:** 10.1101/283556

**Authors:** Isabelle Mifom Vea, Sayumi Tanaka, Tomohiro Tsuji, Takahiro Shiotsuki, Akiya Jouraku, Chieka Minakuchi

**Affiliations:** University of Edinburgh, Institute of Evolutionary Biology, Edinburgh, UK; Nagoya University, Graduate School of Bioagricultural Sciences, Nagoya, Japan; National Agriculture and Food Research Organization, Tsukuba, Japan; Shiname University, Faculty of Life and Environmental Science, Matsue, Japan

**Keywords:** E93, insect metamorphosis, neoteny, hemimetaboly, juvenile hormone, mealybugs

## Abstract

Insect metamorphosis generates reproductive adults and is commonly accompanied by the direct or indirect development of wings. In some winged insects, the imago is altered by life history changes. For instance, in scale insects and mealybugs, reproductive females retain juvenile features and are wingless. The transcription factor *E93* triggers metamorphosis and plays in concert with the juvenile hormone pathway to guarantee the successful transition from juvenile to adult. We previously provided evidence of an atypical down-regulation of the juvenile hormone pathway during female adult development in the Japanese mealybug. Here, we further investigate how *E93* is involved in the production of neotenic wingless females, by identifying its isoforms, assessing their expression patterns and evaluating the effect of exogenous juvenile hormone mimic treatment on *E93*. This study identifies three *E93* isoforms on the 5’ end based on Japanese mealybug cDNA and shows that female development occurs with the near absence of *E93* transcripts, as opposed to male metamorphosis. Additionally, while male development is typically affected by exogenous juvenile hormone mimic treatments, females seem to remain insensitive to the treatment, and up-regulation of the juvenile hormone signaling is not observed. Furthermore, juvenile hormone mimic treatment on female nymphs did not have obvious effect on *E93* transcription, while treatment on male prepupae resulted in decreased *E93* transcripts. In this study, we emphasize the importance of examining cases of atypical metamorphosis as complementary systems to provide a better understanding on the molecular mechanisms underlying insect metamorphosis. For instance, the factors regulating the expression of *E93* are largely unclear. Investigating the regulatory mechanism of *E93* transcription could provide clues towards identifying the factors that induce or suppress *E93* transcription, in turn triggering male adult development or female neoteny.

**Graphical abstract:** 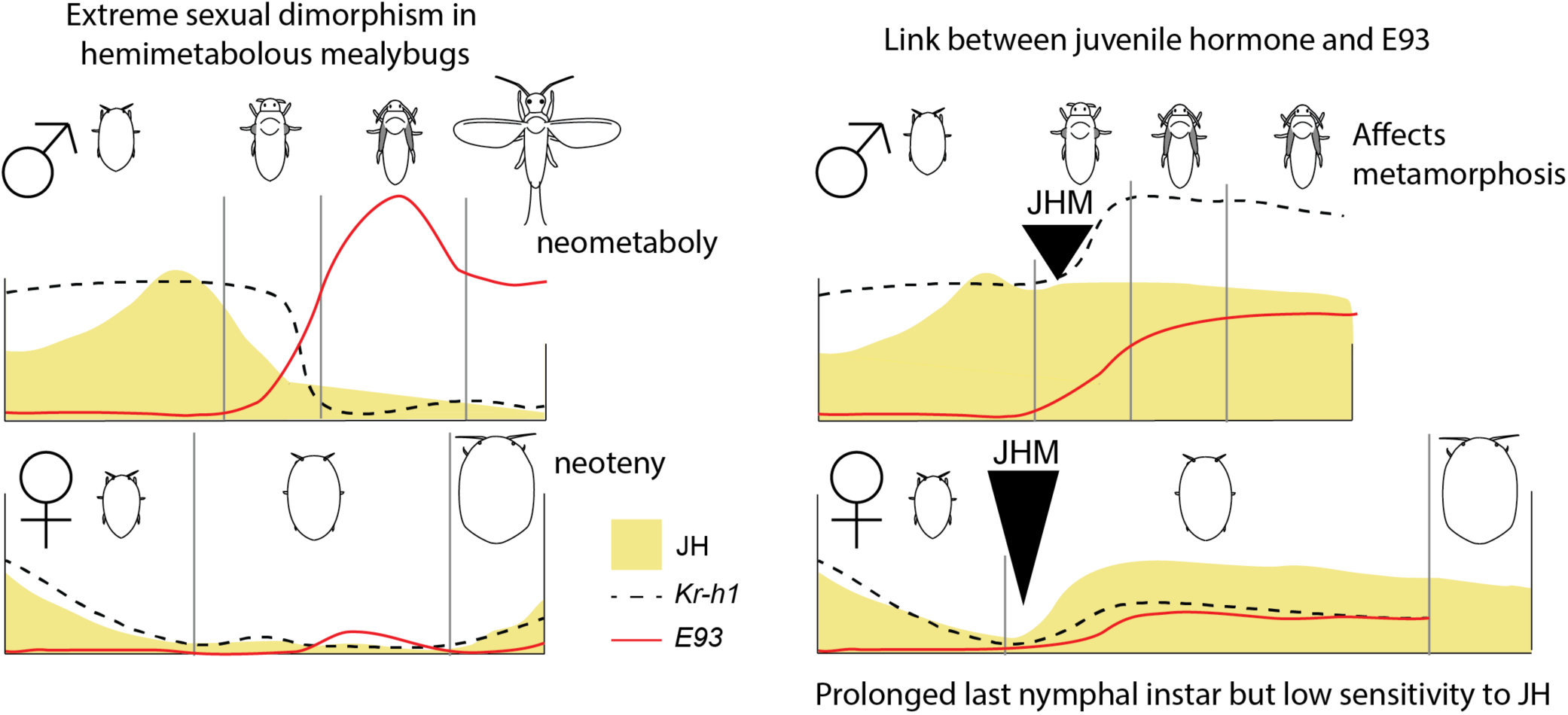

**Highlights:**

- Neotenic female *Planococcus kraunhiae* (Japanese mealybug) develops with low *E93* expression.
- *E93* expression pattern during male development is typical to other insects.
- Juvenile hormone mimic treatment on male prepupae results in decreased *E93* transcripts.
- Juvenile hormone mimic treatment on female nymphs does not have obvious effects on *E93* transcription.
- Female mealybugs have low sensitivity to juvenile hormone mimic treatments compared to males and other insects.

## 1. Introduction

The evolution of metamorphosis undeniably contributed to the diversity of insect forms, life histories and increasing opportunities for ecological niche exploitation (Grimaldi and Engel, 2005; Truman and Riddiford, 1999). While two types of metamorphosis -- holometaboly and hemimetaboly – predominate in insects, in a few lineages, life cycles deviate to form unusual developmental instances, nonetheless leading to important adaptations. For example, hypermetamorphosis, manifested by two types of larvae, arose from holometaboly. Found in Strepsiptera, Meloidae and Rhipiphoridae beetles, neuropteran Mantispidae and many Hymenoptera and some Diptera, hypermetamorphosis is often associated to parasitic life (Truman and Riddiford, 2002) or predatory habits (Belles, 2011). Neometaboly is another metamorphosis found in plant sap-feeding hemimetabolous Paraneoptera (Aleyrodoidea, Thysanoptera, and male Coccomorpha), in which the formation of quiescent stages (prepupae and pupae) is reminiscent to holometaboly (Belles, 2011).

Two hormones orchestrate insect metamorphosis: ecdysone and the juvenile hormone (JH). JH, in particular, dictates the identity of subsequent stages and is therefore of special interest for understanding how peculiar life cycles arise. An essential player, the transcription factor *E93*, triggers adult metamorphosis and is universally up-regulated at the end of insect juvenile development (Ureña et al., 2014). *E93* involvement was first reported in *Drosophila melanogaster* cell death process in the prepupa (Baehrecke and Thummel, 1995; Buszczak et al., 2000; Lee et al., 2000) and acts as a developmental switch to control the responsiveness of target genes during metamorphosis (Mou et al., 2012). Later, functional studies on *E93* orthologs in both hemimetabolous and holometabolous species confirmed *E93* is a universal adult specifier in insect metamorphosis (Ureña et al., 2014). Finally, a communication between *E93* and JH signaling pathway exists. In fact, *Krüppel homolog 1* (*Kr-h1*), an early response gene of JH signaling, acts as a repressor of *E93* until the onset of adult metamorphosis: knocking down *Kr-h1* at the penultimate juvenile instar of hemimetabolous *Blattella germanica* results in the early increase of *E93* (Belles and Santos, 2014). Thereafter, functional studies in hemimetabolous *Cimex lectularius* (Gujar and Palli, 2016) and *Tribolium castaneum* pupal stage (Ureña et al., 2016) confirmed the interaction of JH signaling and *E93* in other insect lineages. Finally, the direct transcriptional repressor role of *Kr-h1* on *E93* promoter region was confirmed in *Bombyx mori* (Kayukawa et al., 2017).

Some unusual life cycle alterations can also result in reproductive forms retaining juvenile features: neoteny. Considered merely as curiosities, neotenic forms originated multiple times in various insects and can be associated to particular adaptive traits. Examples include parasitism in Strepsiptera (Kathirithamby, 2009), Isoptera sociality (Higashi and Abe, 1997; Roisin, 2000), and bioluminescence-generating Elateriformia (fireflies, jewel beetles, click beetles etc…) (Bocakova et al., 2007; South et al., 2011). Nevertheless, few studies have addressed the underlying molecular mechanisms of neoteny. The emergence of diverse ways to metamorphose could be associated to the change in maturation timing (heterochrony), which implies that variations in controlling hormones may be an essential factor in establishing these forms (Gould, 1977). As such, female-specific neotenic forms should be tightly linked to the reproductive function of JH. So far, the main hypothesis for the creation of juvenile-like reproductive females resides in excessive levels of JH, simultaneously affecting female developmental progress and the timing of activating reproductive function (Matsuda, 1976).

Excessive JH titers are indeed observed in termite neotenics of *Reticulitermes speratus*. Here, the female reproductive neotenic caste shows significantly higher JH titers than those of the nymphs or worker castes. Additionally, knocking down JH receptor (*RsMet*) depletes *vitellogenin* transcript levels. However, it is still unclear whether the phenotypic features attributed to neotenics are affected (Saiki et al., 2015). The only other molecular studies on insect neotenic forms were undertaken on holometabolous insects, where the role of ecdysone was investigated as the responsible factor. Strepsiptera (twisted-wing insects) display sex-specific neotenic forms, whereby females in extreme groups are larviform and endoparasitic. The expression patterns of the pupal specifier *broad* in *Xenos vesparum* was also examined. In holometabolous insects, *broad* is up-regulated during the last larval instar, at the onset of metamorphosis with an ecdysone titer increase (Kiss et al., 1988; Konopova and Jindra, 2008; Parthasarathy et al., 2008; Uhlirova et al., 2003). In *X. vesparum*, only males that undergo metamorphosis showed the increase of *broad* BTB domain expression, while it was not observed in last larval instar females (Erezyilmaz et al., 2014). Finally, Cecidomyiidae (gall midges) possess a facultative paedogenetic life cycle where ovaries then differentiate and grow precociously in the larval stage. In this instance, a shift in timing of *ecdysone receptor* and *ultraspiracle* expression kick-starts the facultative life cycle and creates larval reproductive females (Hodin and Riddiford, 2000).

Scale insects and mealybugs (Coccomorpha) belong to Hemiptera, an insect order that mostly develop through hemimetaboly. However, Coccomorpha species have departed from the traditional nymphal instars with progressive wing growth. Males undergo two quiescent stages reminiscent to complete metamorphosis (neometaboly, as mentioned above). In striking contrast, reproductive females retain juvenile features, as they develop through successive molts without wing growth, reduction of nymphal stages and features linked to mobility in many species (Gullan and Kosztarab, 1997). This life history trait not only gives rise to extremely sexually dimorphic organisms, but offers a successful strategy, as plant-sap feeding insects, to allocate energy to reproduction, sedentarize and adapt to their host-plant habitats. Female scale insects and mealybugs are often described as neotenic (Danzig, 1980; Koteja, 1990). However, which type of neoteny, or the mechanisms by which adult females keep juvenile features remains unknown. This prevents us from understanding the link between the development and evolution of neoteny. More importantly, this lineage includes some of the most damaging agricultural pests in human activities, likely a consequence of the evolution of neotenic females and their adaptive life history to host plants.

We previously presented a study on the variation of JH in the Japanese mealybug *Planococcus kraunhiae* (Pseudococcidae) to examine variations between male and female development as a possible mechanism leading to extreme sexual dimorphism in scale insects. In addition to significant differences observed in JH early-response gene *Kr-h1* when male and female mealybugs start to differentiate, we reported that JH signaling remained unusually low throughout female adult development, suggesting a different JH regulation of female neoteny in this case (Vea et al., 2016). To further examine the involvement of *E93* in female neoteny, in relation to the JH signaling, we compared the expression pattern of *E93*, in *P. kraunhiae* male and female postembryonic development and performed hormonal assays using a JH mimic, pyriproxyfen. We hypothesize that in scale insects, females fail to express *E93*, resulting in the maintenance of juvenile features in their external morphology even after reproductive maturation. Additionally, we test whether increasing levels of JH during the last nymphal instars in females readjusts the expression of *Kr-h1* at similar levels as seen in males, which in turn could allow to initiate *E93* expression. It turned out that females are insensitive to exogenous JHM treatment in the context of the effects on *Kr-h1* and *E93* expression as well as on adult development.

## 2. Materials and methods

### 2.1. Mealybug rearing and sampling strategy

Mealybug culture and sampling strategy for gene expression profile are described in a previous study on JH variations in the *P. kraunhiae* (Vea et al., 2016). In this study, we carried out an independent sampling to ensure reproducibility of the previous study. As such, we collected samples every 24 hours after oviposition. Eggs oviposited during the first day were used for male-biased samples and eggs oviposited during the fifth day for female-biased samples (see Vea et al., 2016 for sex-biased sample strategy). All stages are abbreviated as follows: E = embryonic stage after oviposition, N1= first-instar nymph, N2= second-instar nymph, N2f= female second-instar nymph, N2m= male second-instar nymph, N3= female third-instar nymph, pre=male prepupa, pu= male pupa, m=male adult, f=female adult.

### 2.2. cDNA cloning and identification of sequences

The total RNA of pooled individuals from different stages was extracted with TRIzol (as described in Vea et al., 2016) and Oligo-dT-primed reverse transcription was performed with the PrimeScript II 1^st^ strand cDNA synthesis kit (Takara Bio, Shiga, Japan). The conserved region of *E93* sequence in *B. germanica* was blasted against a transcriptome of *P. kraunhiae* [accession number DRA004114; (Sugahara et al., 2015)], and primers for RT-PCR were designed to amplify a partial region of the gene. Primers for RACE PCR were designed based on this partial sequence, and 5’ and 3’ RACE was conducted with SMARTer RACE cDNA Amplification Kit (Takara Bio USA, Inc., Mountain View, CA) in order to retrieve the full-length cDNA sequences. All PCR products were cloned in a pGEM-T Easy Vector (Promega, Madison, WI) and sequenced. DNA sequence data were deposited in the DDBJ/EMBL-Bank/GenBank International Nucleotide Sequence Database with the following accession numbers: *PkE93* isoform 1 (LC374380), *PkE93* isoform 2 (LC374381) and *PkE93* isoform 3 (LC374382). The primer sequences are listed in Table S1.

### 2.3. RNA extraction and quantitative RT-PCR

Total RNA was extracted from all samples using the sex-biased sampling as described in Vea et al. (2016). Each sample consisted of 0.5 to 2 mg of pooled individuals homogenized in TRIzol reagent, total RNA was extracted using nuclease-free glycogen (Thermo Fisher Scientific) as a carrier, and reverse transcribed using the PrimeScript RT reagent Kit with the gDNA Eraser (Takara Bio). Expression profiles for the post-oviposition development of males and females were established by quantifying the levels of transcripts for targeted fragments using absolute quantitative RT-PCR (qRT-PCR), performed on a Thermal Cycler Dice Real Time System (model TP800, Takara Bio) as described previously (Vea et al., 2016). Six serial dilutions of a plasmid containing a fragment of each gene was used as the standard. Primer sequences for each *PkE93* isoform used for qRT-PCR are listed in Table S1. The values obtained by the second derivative maximum (SDM) method were normalised with the *ribosomal protein L32* (*rpL32*) transcript levels. Primers for *PkrpL32*, our reference gene, were from our previous study (Vea et al., 2016).

### 2.4. JH mimic assays on male prepupae and female juvenile instars

For JH mimic (JHM) treatments, we applied 2 µL of pyriproxyfen (5 mM dissolved in methanol) to batches of 3 to 5 male prepupae on a filter paper, 24-48 hours after molting. Excess chemical solution was immediately absorbed by the filter paper. After treatment, we waited for the solvent to evaporate completely before transferring them in 1.5 mL microcentrifuge tubes (Ina Optica, Osaka, Japan), bearing a paper disc (8 mm in diameter, 1.5 mm in thickness; Toyo Roshi, Japan) with 10 µL of distilled water to control humidity. Female N3D0 (0-24 h after molting to the N3 stage) were treated in the same manner except that 0.5 µL of pyriproxyfen (20 mM dissolved in methanol) was applied on the tergite of individual N3D0. After the N3D0 started to move again, it was transferred into a glass dish containing a sprouted broad bean on top of a filter paper. The glass dishes were sealed with parafilm. Treated samples were left to incubate at 23°C for various numbers of days after treatment before being homogenized in TRIzol and stored at −80 °C for RNA extraction. RNA extraction, reverse transcription and qRT-PCR analyses were performed as described in 2.3.

### 2.5. Graphs and statistical analyses

Graphs from qRT-PCR SDM values were generated using the R package ggplot2 and the statistical significance of JHM effects were calculated by comparing the overall effect of the mimic over the days using a linear model. We also tested the effect of JHM on individual days 4 and 8 after treatment in females using the Student’s *t*-test. All analyses using qRT-PCR SDM values are detailed and publically available in GitHub and Zenodo (https://zenodo.org/badge/latestdoi/116843862).

## 3. Results and Discussion

### 3.1. Structure of E93 in P. kraunhiae

The *PkE93* sequence identified from RT-PCR includes a Pipsqueak DNA binding domain characteristic of *E93* (Siegmund and Lehmann, 2002) and highly conserved (**Fig. 1A**). Using 5’ and 3’ RACE PCR combined to designed primers from the conserved region, we cloned and sequenced three complete transcripts of 5008, 5056 and 5233 bp long. All of them resulted from either usage of different transcription initiation site and/or alternative splicing on 5’end only (**Fig. 1B**). Each isoform was arbitrarily designated as *PkE93-1, PkE93-2* and *PkE93-3*. The region common to all transcripts counts 4676 bp, *PkE93-1* and *PkE93-3* predicted protein sequence is of 1050 aa, while *PkE93-2* has a predicted protein sequence of 1090 aa. In summary, *PkE93-1* and *PkE93-3* differ in the 5’ untranslated region, while *PkE93-2* has a longer coding region.

**Figure 1:**
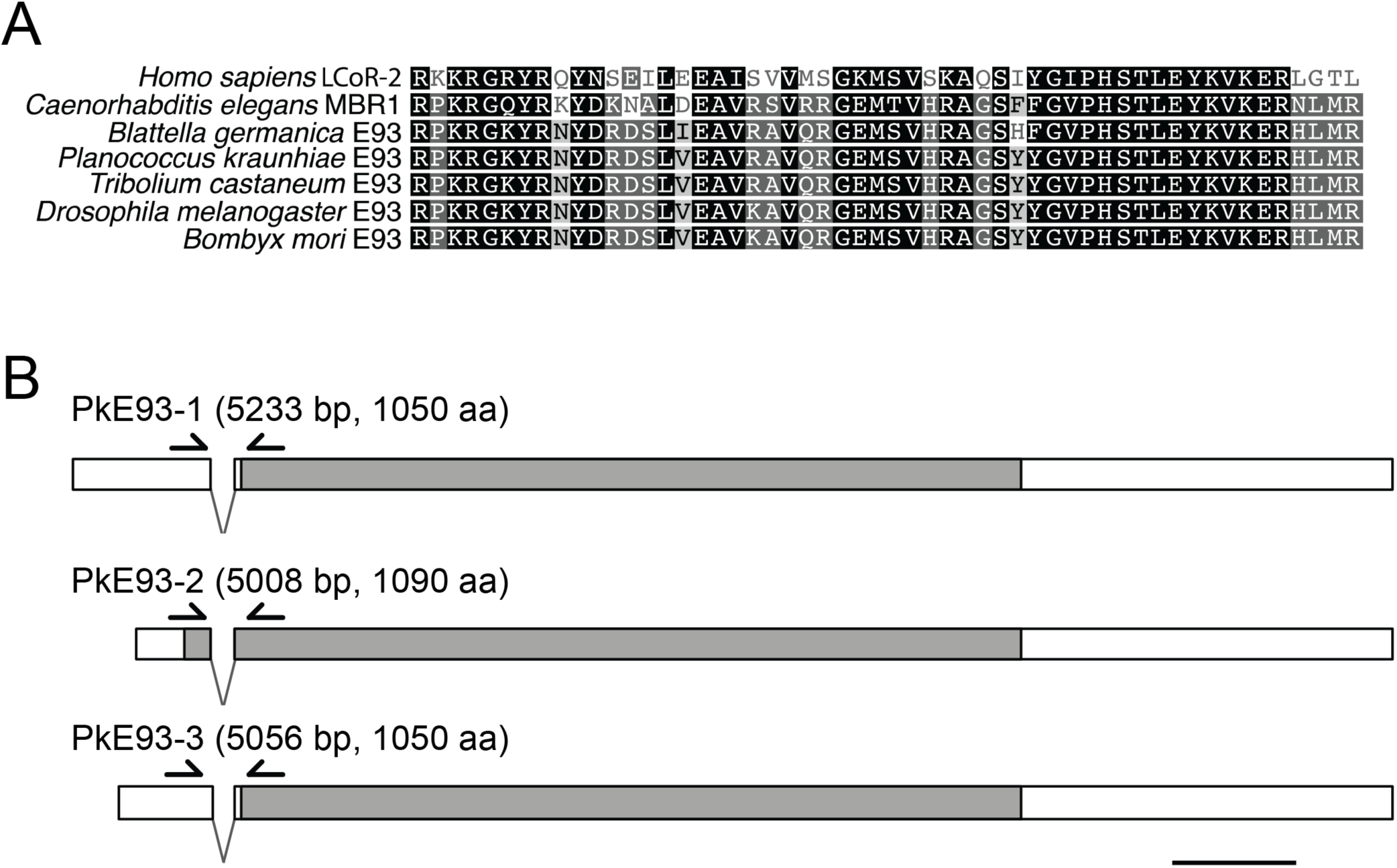
Identification of PkE93 sequence and the structure of its isoforms. A. Amino acid alignment of the Pipsqueak DNA-binding domain with other insects’ E93, *Homo sapiens* LCoR-2 and *Caenorharbidtis elegans* MBR1 [sequences from (Ureña et al., 2014)]. B. General structure of cDNA sequences obtained from 5’ and 3’ RACE PCR. The sequences are identical on the 3’ end, while the 5’ end differ among the three isoforms, the common region is of 4676 bp starting from 3’ end. Grey: open reading frame. Arrows: primers designed for qRT-PCR. Scale bar: 500 bp.

### 3.2. Sex-specific expression profiles of E93

We first examined the expression profile of each *PkE93* isoform during the post-oviposition development in male and female mealybugs, using absolute qRT-PCR (**Fig. 2A**). *PkE93-1* isoform has the highest expression compared to the two other isoforms. Generally, *PkE93-1* stays at low levels in the embryo, N1 and N2 in both males and females (**Fig. 2A; top**). In males only, *PkE93-1* suddenly increases at the beginning of prepupa and reaches a peak of expression before adult metamorphosis. In females, however, *PkE93-1* does not show such dramatic expression, although slight increases at the end of N2 and N3 are observed (**Fig. 2B**). *PkE93-2* isoform followed a similar expression pattern although around 20-fold lower (**Fig. 2A; middle**), which suggests that *PkE93-2* may be a minor isoform. Finally, *PkE93-3*, despite its also lower expression, shows a distinct pattern during the embryonic stage (**Fig. 2A; bottom**), with peaks of expression in both males and females, similar to that of *PkKr-h1* (**Fig. S1**). At the end of N2, *PkE93-3* expression also increases in males but decreases during the pupal stage, while its expression reaches near-zero levels after N2 in females. *PkE93* expression differs between males and females more strikingly than the early JH-response gene *PkKr-h1*. This is especially true towards the end of post-embryonic development, where a peak of expression is observed in males, but not in females. At this point, males enter quiescent stages (prepupa and pupa) where wings develop. In contrast, females molt only once then become reproductively mature but retain juvenile features. Along with previous results on *PkKr-h1* and *Pkbr* (Vea et al., 2016), we suggest that sex-specific expression patterns of *PkE93* may contribute to the development of sexual dimorphism in mealybugs.

**Figure 2:**
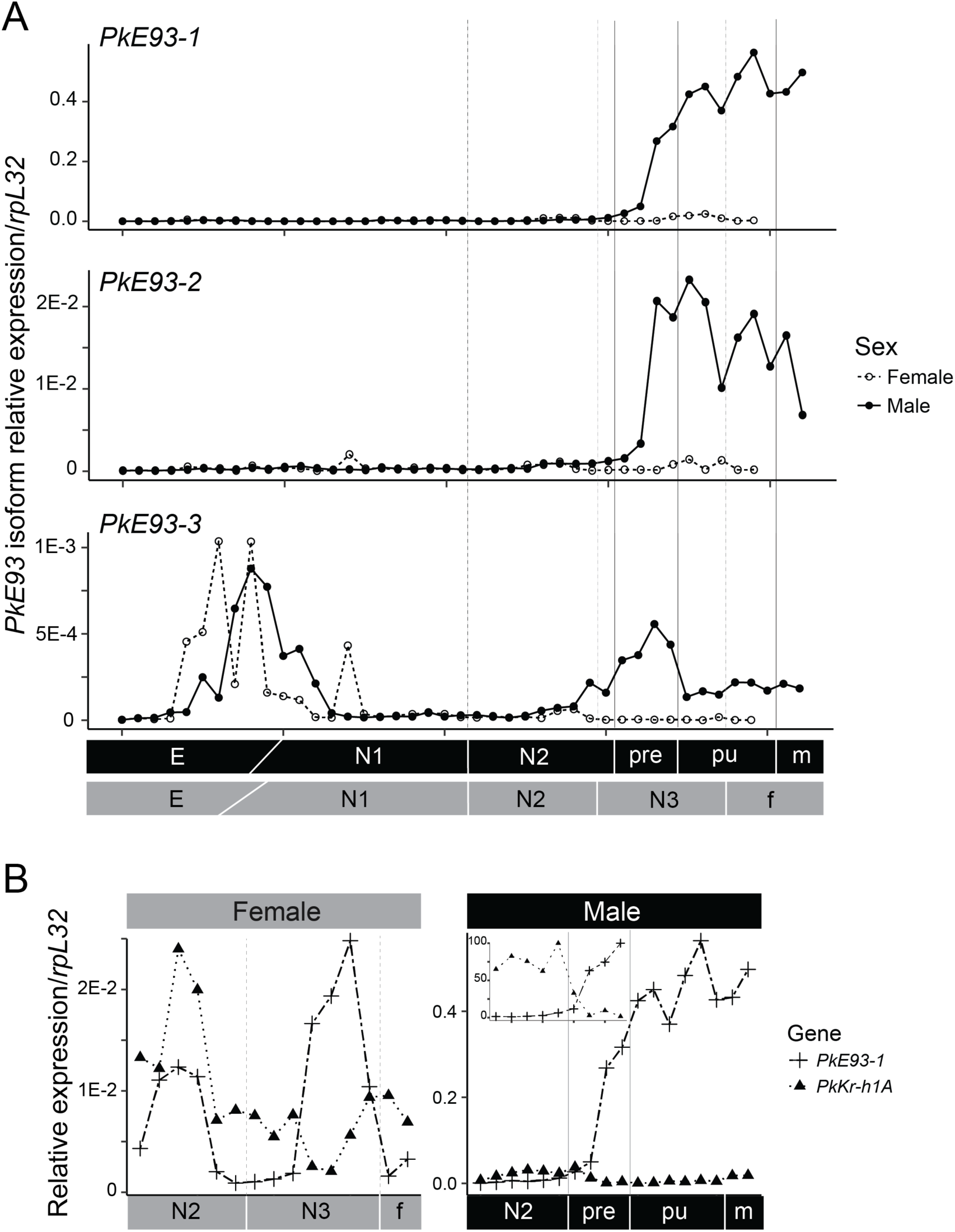
Expression profiles of *PkE93* isoforms throughout post-oviposition in males and females, and comparison with those of *PkKr-h1* at the end of development. A. *PkE93-1, PkE93-2* and *PkE93-3* expression profiles from qRT-PCR of samples collected every 24 hours from oviposition. The primers used for qRT-PCR are shown in Fig. 1 and their sequences are listed in Table S1. B. Comparison of *PkE93-1* and *PkKr-h1A* expression from the second-instar nymph, when female and male can be differentiated, to the adult stage. The small graph on top left part of male expression graph shows the relative expression in percentage when the switch of expression occurs between the two genes.

### 3.3. Expression pattern of male mealybugs consistent with JH signaling of other insects

The origin of holometaboly has been debated for decades. Although the consensus is that endocrinological changes are responsible for the transition between hemimetaboly and holometaboly, two hypotheses were advanced regarding the details of stage homologies [see details in (Truman and Riddiford, 1999)]. More recently, based on study of *E93* expression in *B. germanica*, it was proposed that the last nymphal stage in hemimetabolous insects is ontogenetically homologous to the pupal stage of holometabolous insects because of the similar *E93* expression pattern during these stages (Belles and Santos, 2014). Furthermore, the propupal and pupal stages in neometabolous Thysanoptera, could be together homologous to the holometabolous pupal stage (Minakuchi et al., 2011). In scale insects, a similar neometabolous development occurs in males (stages traditionally coined as “prepupa” and “pupa”) and our expression profile pattern in male *P. kraunhiae* (**Fig. 2A**) shows that *PkE93* expression starts at the beginning of the prepupa and peaks at the beginning of the pupal stage. Finally, hormonal treatment using JHM at the beginning of the prepupal stage, when *Kr-h1* expression drops suddenly, prevents adult metamorphosis by creating a supernumerary pupal stage (Vea et al., 2016). In addition to maintaining high levels of *PkKr-h1* for at least six days, JHM treatment results in the significant decrease of *PkE93-1* and *PkE93-2*, while *PkE93-3* is not significantly affected (**Fig. 3A**). Although functional analyses are still necessary to confirm this in mealybugs, the effects of JHM on *E93* expression in neometabolous males are congruent with previous results found in other holometabolous and hemimetabolous insects, and *Kr-h1* and *E93* must be acting as reciprocal inhibitors.

**Figure 3:**
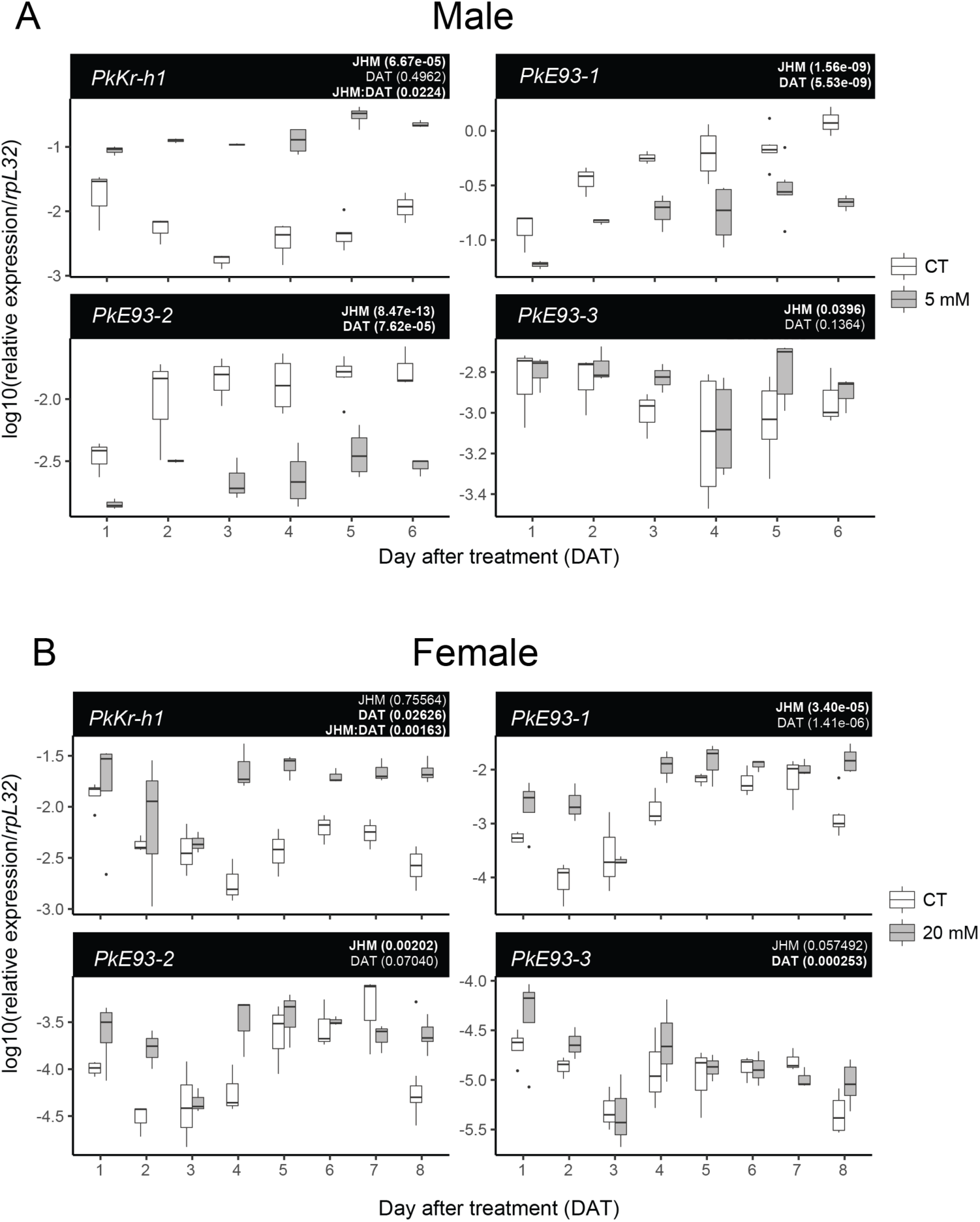
Effect of juvenile hormone mimic treatment on *Kr-h1* and *E93* expression at the end of male and female development. The transcript levels were analyzed with qRT-PCR, and normalized with the *rpL32* levels. A. Expression of *PkKr-h1, PkE93-1, PkE93-2* and *PkE93-3*, 1 to 6 days after male 1 day old prepupae (PreD1) were treated with 5 mM pyriproxyfen or methanol. B. Expression of *PkKr-h1, PkE93-1, PkE93-2* and *PkE93-3*, 1 to 8 days after 0-24 h old third instar females (N3D0) were treated with 20 mM pyriproxyfen or methanol. Statistical significance: we used a linear model in R on log10 transformed data, to test the significance of pyriproxyfen treatments and also considered day after treatment as a factor (see GitHub link in Materials and Methods for details on the analysis). P-values for each factor are indicated on top right side of each graph. We first considered an interaction between the two tested variables in our linear model. If the interaction is significant, the p-values of treatment (JHM), day after treatment (DAT) and interaction between them (JHM:DAT) are indicated (as it is the case for *PkKr-h1*). If no interaction was found, we used the linear model excluding the interaction and indicated p-values for treatment and day after treatment only (for all three *PkE93* isoforms). We considered an effect significant when p-value < 0.01.

### 3.4. Female atypical regulation of JH signaling and E93

Based on our expression profile of *PkE93* in females, all isoforms are very low throughout the successive molting events, contrasting with male neometabolous development. However, if we compare *PkE93* and *PkKr-h1* expression only during the last instar nymphs, we see that even at very low levels, a slight decrease of *PkKr-h1* is accompanied with a small peak of expression of *PkE93-1* (**Fig. 2B**). Although the female expression is 20-fold lower than in males, this small peak could explain the sexual maturation of females after N3, while somatic differentiation does not occur in the imago. Additionally, this small peak likely indicates tissue-specific expression, like ovaries and related reproductive organs. As *PkKr-h1* expression begins to decrease progressively at the penultimate nymphal stage in females (**Figs. 2B and S1**; Vea et al. 2016), we decided to increase the expression of *Kr-h1* with JHM treatment at the early last nymphal stage (N3D0). By doing so, we assessed whether *Kr-h1* can artificially reach a threshold necessary for the switch to adult fate controlled by *E93*. Twenty mM of pyriproxyfen were applied on N3D0, concentration four times higher than male treatments. Following this treatment, 7 out of 19 (37 %) treated individuals died before the last molt (**Table 1**). The 12 survivors (63 %), showed a tendency to prolong the N3 stage. While the control samples molted 9.6±1.5 days (N=14) after treatment, 12 out of 19 JHM-treated individuals molted to adult stage 13±2.5 days after treatment. We also followed the effect of JHM treatment on *PkKr-h1* and *PkE93* expression (**Fig. 3B**), one to eight days after treatment (last days before the female adult molt). At 20 mM, pyriproxyfen induced the up-regulation of *PkKr-h1* after four days, which then lasted several days. However, it is worth mentioning that *PkKr-h1* levels were significantly lower than the response observed in males, even with a concentration of pyriproxyfen four times lower. We suggest that female mealybugs may have mechanisms preventing them to respond to JH as sensitively or efficiently as in males. Previously, we showed that *PkMet* and *PkTai*, forming the JH receptor complex, were highly expressed at the end of male development, while in females the expression remained low (Vea et al., 2016). After JHM treatment on females at N3D0, although *PkKr-h1* starts to be affected only four days after treatment by an upregulation, the expression of all *PkE93* isoforms does not change significantly over time (interaction between treatment and time not significant; **Fig. 3B**). However, when removing the interaction, the treatment alone leads to an overall significant effect for two isoforms; this is probably explained by the increase in expression at Day 2, Day 4 and Day 8 after treatment for *PkE93-1* and *PkE93-2*. We therefore conclude that applying JHM on early ultimate juvenile instars in females does not significantly change *PkE93* expression overtime, but a small effect might be observed at Day 8 after treatment by an increase of *PkE93-1* and *PkE93-2*.

**Table 1:**
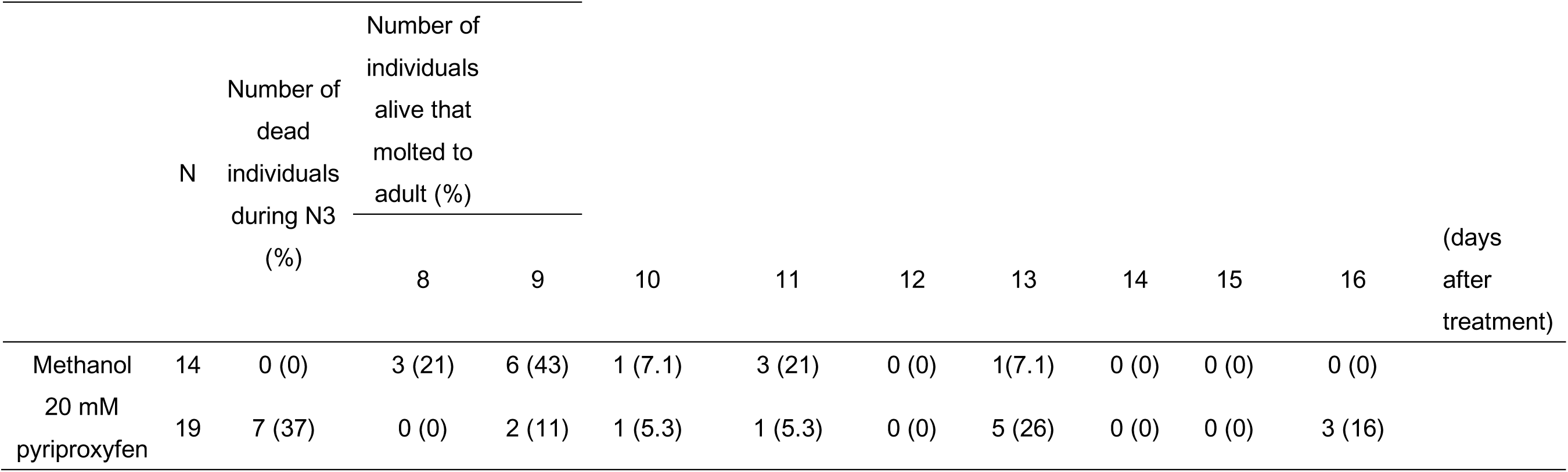
Effect of pyriproxyfen treatment on last nymphal instar females on adult metamorphosis

Several studies have demonstrated that at the onset of metamorphosis in holometabolous insects, juvenile instars need to reach a critical weight for adult metamorphosis (Davidowitz et al., 2003; Nijhout, 2003; Nijhout et al., 2014). In mealybugs, males and females are phenotypically identical before their development starts to diverge. If we consider that critical weight is a factor in at least male metamorphosis, the threshold should be reached at the start of prepupa, where we see the sudden decrease of *PkKr-h1*. By contrast, females should attain their critical weight for metamorphosis in N3 stage. In females, *PkKr-h1* down-regulation takes place in a progressive decrease from the middle of N2 rather than the sudden decline of transcripts. Moreover, *Pkjhamt* (*juvenile hormone acid O-methyltransferase*), involved in the last steps of JH biosynthesis, starts decreasing even at the beginning of N2, while in males, the transcripts are still expressed until prepupa (Vea et al., 2016). This indicates an early arrest in JH synthesis in female development. Another possibility is that a sudden decrease combined with attaining a threshold of *Kr-h1* expression is necessary to induce *E93* expression. In this case, maintaining low levels of *PkKr-h1* during female development may explain why *PkE93* expression never peaks.

## 4. Concluding remarks

JH anti-metamorphic role in insect metamorphosis suggests that high levels of this hormone delays metamorphosis. Female neoteny has therefore been believed to be the result of disrupted JH down-regulation leading to constantly high JH titers. Although this hypothesis was first proposed based only on the differential size of scale insect *corpora allata* (Matsuda, 1976), we recently reported contradictory evidence that the last juvenile stages in female Japanese mealybug develop under surprisingly lower JH titers compared with males (Vea et al., 2016). In this study, we provide additional data to support an atypical hormonal regulation in mealybugs, and show the first example of possible failure in *E93* induction, linked to neotenic reproductive females in an hemimetabolous insect. The response of female development to JH modulations is intriguing and suggests that mealybug neotenic forms are insensitive to JH signaling as opposed to males, has consequences on *E93* expression, and leads to extreme sexual dimorphism. So far, gene expression manipulation on mealybug juvenile stages by dsRNA injection has been ineffective. Alternatively, identifying suppressors of *E93* promoter in neotenic females, coupled with functional studies through genome editing should provide novel insights on the function and interaction between *E93* and JH signaling pathway in neotenic female scale insects.

## Acknowledgements

We thank Jun Tabata for providing the Japanese mealybugs, Ken Miura and Xavier Belles for the valuable comments on this study. All the work presented here was conducted at Nagoya University and was funded by a JSPS postdoctoral fellowship for overseas researchers to IMV and a Grant-in-aid for Scientific Research (15K07791) to CM from JSPS.

## List of supplementary material

Table S1: List of primers used for cloning, 5’ and 3’ RACE PCR and quantitative RT-PCR.

Figure S1: Expression profile of *PkKr-h1* during male and female Japanese mealybug development after oviposition

GitHub/Zenodo repository for data analysis: https://zenodo.org/badge/latestdoi/116843862 Protocol repository: protocols for sex biased collecting strategy in mealybugs (dx.doi.org/10.17504/protocols.io.mh9c396) and pyriproxyfen treatments on mealybugs (dx.doi.org/10.17504/protocols.io.miac4ae)

