## Supplementary Materials for "E93 expression and links to the juvenile hormone in hemipteran mealybugs with insights on female neoteny"

### Supplementary material

Table S1. List of primers used for cloning, 5’ and 3’ RACE PCR and quantitative RT-PCR

| **Primer name** | **Sequence 5' -> 3'** | **Used for** | **Expected size** |
|---|---|---|---|
| PkE93-F1 | CAGCAACAACAACAACAACAGC | cloning | 1864 |
| PkE93-R1 | TATCAACGCTGGCAATTTGAGA | cloning | -- |
| PkE93-RR1 | GGCTGGTGATGGGAACTTTGGACTGA | 5'RACE | -- |
| PkE93-RR2 | TGGATTTGCATGCCCTATCAGAGACG | 5'RACE | -- |
| PkE93-RR3 | GCTGGTAATGCTGCCATTTCCTTCTGC | 5'RACE | -- |
| PkE93-RF1 | GCTATCCTCCGTTGTCGCCAAAGACAC | 3'RACE | -- |
| PkE93-RF2 | GCCTCAGCTTCAGCTGCCAAGAGTGT | 3'RACE | -- |
| PkE93-QF1 | TCATCACCATTGCCTATGAACC | qRT-PCR | 115 (PkE93-QR1 as the reverse primer) |
| PkE93-QR1 | TCATGTGTGACAATGGCAAGTC | qRT-PCR | -- |
| PkE93-1-QF1 | TCAAACTGTTTCGATGTGAGTAAGG | qRT-PCR | 105 (PkE93-1_2_3-QR1 as reverse primer) |
| PkE93-2-QF1 | CCGAACGTTACGGTGTGATTT | qRT-PCR | 145 (PkE93-1_2_3-QR1 as reverse primer) |
| PkE93-3-QF1 | CGCAGCATTTGATCCAAAAA | qRT-PCR | 136 (PkE93-1_2_3-QR1 as reverse primer) |
| PkE93-1_2_3-QR1 | TGCTGGTAATGCTGCCATTT | qRT-PCR | -- |

Figure S1: Expression profile of *PkKr-h1* during male and female Japanese mealybug development after oviposition

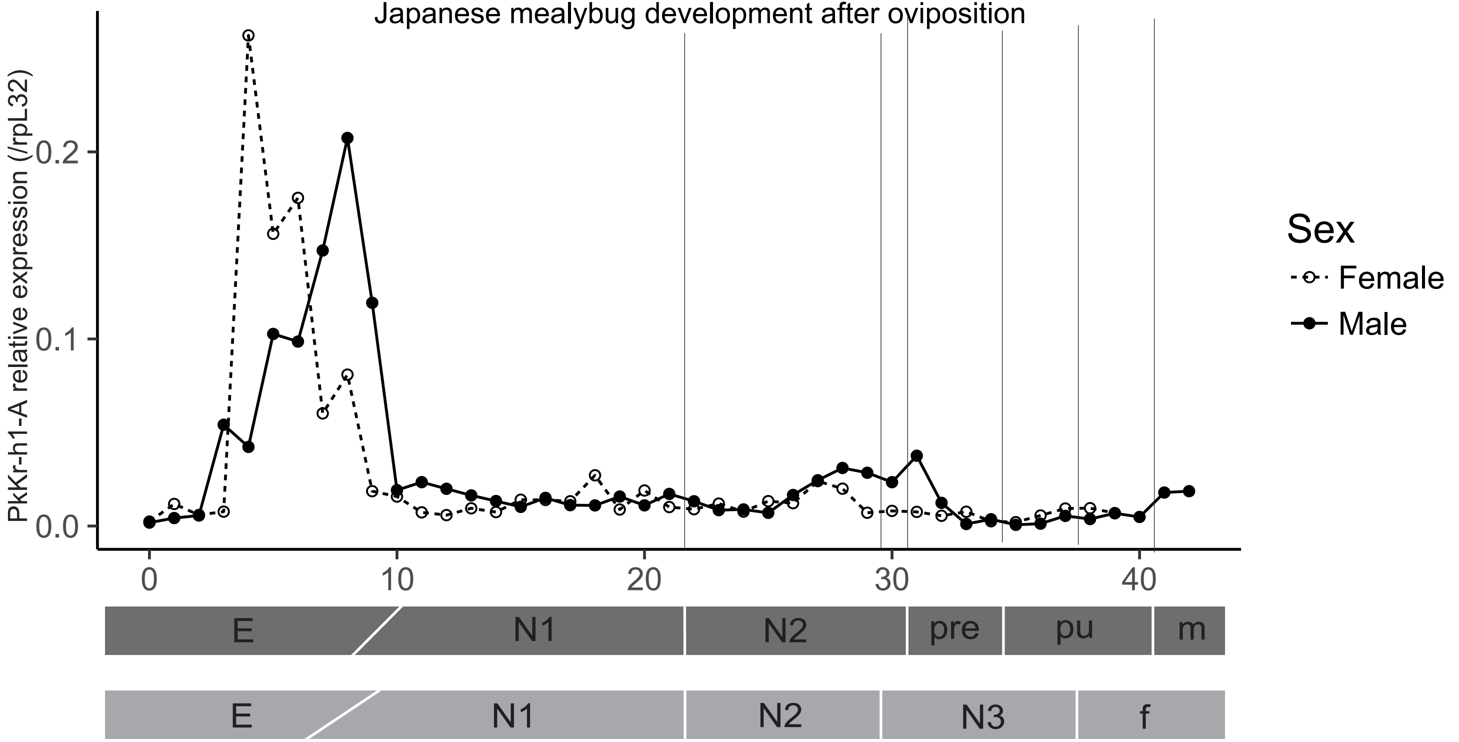
